# An Exon Skipping Screen Identifies Antitumor Drugs That Are Potent Modulators of Pre-mRNA Splicing, Suggesting New Therapeutic Applications

**DOI:** 10.1101/584441

**Authors:** Yihui Shi, Walter Bray, Alexander J. Smith, Wei Zhou, Joy Calaoagan, Chandraiah Lagisetti, Lidia Sambucetti, Phillip Crews, R. Scott Lokey, Thomas R. Webb

**Affiliations:** Bioscience Division, SRI International, Menlo Park, CA 94025; Department of Chemistry and Biochemistry, University of California, Santa Cruz, Santa Cruz, CA 95064

**Keywords:** Spliceosome, RNA processing, high-throughput screening, modulators of pre-mRNA splicing, kinase inhibitors, cancer

## Abstract

Agents that modulate pre-mRNA splicing are of interest in multiple therapeutic areas, including cancer. We report our recent screening results with the application of a cell-based Triple Exon Skipping Luciferase Reporter (TESLR) using a library that is composed of FDA approved drugs, clinical compounds, and mechanistically characterized tool compounds. Confirmatory assays showed that three clinical antitumor therapeutic candidates (milciclib, PF-3758309 and PF-030871) are potent splicing modulators and that these drugs are, in fact, nanomolar inhibitors of multiple kinases involved in the regulation the spliceosome. We also report the identification of new SF3B1 antagonists (sudemycinol C and E) and show that these antagonists can be used to develop a displacement assay for SF3B1 small molecule ligands. These results further supports the broad potential for the development of agents that target the spliceosome for the treatment of cancer and other diseases, as well as new avenues for chemotherapeutic discovery.

## Introduction

The use of targeted high-throughput screening (HTS) of recently available compound libraries composed of drugs, clinical compounds and advanced tool compounds offers the biomedical research community the opportunity to elucidate the mechanism of action (MOA), on-target specificity and potential for clinical repositioning of specific drugs, while at the same time developing a refined drug candidate profile for researchers in specific areas of drug discovery and drug development. The spliceosome is accountable for the post-transcriptional processing of pre-mRNA in the cells of metazoans by catalyzing the regulated exclusion of intervening sequences (introns) and the ligation of coding regions (exons) to produce mature mRNAs, and has recently emerged as a novel target in several therapeutic areas.(Wahl, Will et al. 2009) Small molecules that affect AS have been of interest for numerous therapeutic applications since they impact cellular function by modifying the abundance of different splicing isoforms that play a role in numerous disease states.(Ohe and Hagiwara 2015)

Given the important role that the spliceosome plays in the determination of cellular and organismal phenotypes it is not surprising that the function of the spliceosome is aberrant in most tumors.(Chen and Weiss 2015) Indeed, numerous genes are subject to splicing events that can be either oncogenic or serve to limit potential tumorigenesis, examples of this include BCL-X, VEGF-A, FAS, PKM or MDM2.(Bonnal, Vigevani et al. 2012) Additionally, numerous recurrent mutations occur in spliceosome regulatory components (including SF3B1, SRSF2, U2AF1 and others) in the myelodysplastic syndromes and other cancers.(Graubert, Shen et al. 2012) These mutations result in a ‘change in function’ of the mutant spliceosome and a consequential change in the AS profile in the cells expressing these mutant proteins.(Brooks, Choi et al. 2014, Dvinge, Kim et al. 2016, Obeng, Chappell et al. 2016)

In parallel to these recent discoveries, there has been a proportional upsurge in interest in the potential application of several recently discovered small molecule modulators of pre-mRNA splicing to cancer chemotherapy.(Lee and Abdel-Wahab 2016, Bates, Morris et al. 2017, Leon, Kashyap et al. 2017) This effort has resulted in two Phase I clinical studies, and advanced pre-clinical development, for a series of ligands of the SF3B1 spliceosomal protein. These innovative drugs include a derivative of the natural product pladienolide (E7107),(Kotake, Sagane et al. 2007) a synthetic analog of pladienolide(Mizui, Sakai et al. 2004, Sakai, Sameshima et al. 2004) (H3B-8800),(Seiler, Yoshimi et al. 2018) and sudemycin D6 (SD6)(Lagisetti, Palacios et al. 2013) a simplified synthetic analog of a natural product (FR-901,464).(Kaida, Motoyoshi et al. 2007) SD6 is currently actively advancing through the ‘investigational new drug’ (IND) development process. Although the natural products which inspired this class of drugs were initially described as “splicing inhibitors”,(Kaida, Motoyoshi et al. 2007, Kotake, Sagane et al. 2007) we now know that SF3B1 targeted agents act as potent modulators of AS through a change in 3’ splice-site fidelity.(Corrionero, Minana et al. 2011, Fan, Lagisetti et al. 2011, Wu, Fan et al. 2018) Tumor cells exposed to the splicing modulatory natural products (and analogs) display a profound change in AS,(Fan, Lagisetti et al. 2011, Wu, Fan et al. 2018) which is similar to the pharmacology that has been observed with kinase inhibitors that interfere with the regulatory phosphorylation of splicing factors.(Bates, Morris et al. 2017)

Although the complete MOA for the tumor selective toxicity of these agents remains to be fully elucidated, several mechanism types have been delineated. An early mechanism to be recognized was the sensitivity of tumor cells bearing spliceosomal mutations, such as CLL carrying SF3B1 mutations, (Xargay-Torrent, Lopez-Guerra et al. 2015) and MDS carrying U2AF1 mutations.(Shirai, White et al. 2017) Additionally, it was found that tumors driven by MYC(Hsu, Simon et al. 2015) or KRAS(Fraile, Manchado et al. 2017) are also sensitized to this class of drugs. More recently proposals have appeared for two additional general mechanisms that may account for the observed selective action of SF3B1 targeted agents in certain cancers, the first proposes that ~11% of all cancers have a partial copy of wild-type SF3B1 protein, which renders these tumors sensitive to SF3B1 targeted drugs;(Paolella, Gibson et al. 2017) another recent publication presents data which is consistent with the idea that certain tumors driven by BCL2A1, BCL2L2 and MCL1 are especially susceptible to SF3B1 targeted agents.(Aird, Teng et al. 2019) It is certainly possible that multiple mechanisms can account for the selective tumor toxicity that has been observed with these agents, which supports the concept that these agents have good potential for broad application in cancer chemotherapy.(Lee and Abdel-Wahab 2016)

Given these new insights into the relationships between carcinogenesis and spliceosome function we initiated a project aimed at the discovery of additional small molecules that target the spliceosome. This has been facilitated by our Triple-Exon Skipping Luciferase Reporter (TESLR) cell-based HTS assay,(Shi, Joyner et al. 2015) which reports on a particular type of triple-exon skipping event in MDM2 pre-mRNA that we first observed in tumor cells treated with sudemycin analogs.(Fan, Lagisetti et al. 2011) We applied this HTS assay to the unbiased screen of a collection of all FDA approved drugs, bioactive compound collections, and compounds in Phase I through Phase III clinical trials, in order to build on our previously reported pilot screening results with this assay, which identified two known cyclin-dependent kinases (CDK) inhibitors that were found to also inhibit several members of the cdc-like kinase (CLK) family. The hits from this pilot screen were then developed into SRI-30125, a selective inhibitor of CLKs 1, 2, and 4.(Shi, Park et al. 2017) Importantly this work also showed that aminopurvalanol,(Chang, Gray et al. 1999) a commonly used tool compound that is often considered a “selective CDK2 inhibitor” is actually a multi-kinase inhibitor and a potent modulator of AS, which we showed acts through the inhibition of CLKs 1, 2, and 4.(Shi, Park et al. 2017)

This paper describes the results from our clinical compound focused HTS screen (see Figure 1), which led to the identification of three potent post-Phase I splicing modulatory drugs (**1**, **2**, and **3**: see Table 1),(Roberts, Ung et al. 2008, Brasca, Amboldi et al. 2009, Murray, Guo et al. 2010) that have not been previously recognized as splicing modulators. These investigational drugs were developed to target oncogenic kinases and were not previously reported to inhibit splicing regulatory kinases. All of these confirmed hits have been previously reported to display potent *in vivo* antitumor activity, which was accounted for due to their activity to other kinases (CDK2, PAK4 or FAK, respectively), however we now report that they are also nanomolar inhibitors of subsets of the multiple kinases involved in the regulation of spliceosome activity. These observations also align with the previous report that another clinical antitumor agent CX-4945 (Silmitasertib) inhibits splicing regulatory kinases and modulates alternative splicing.(Kim, Choi et al. 2014) CX-4945 was developed as a CK2 inhibitor but was later recognized to be a modulator of alternative splicing following the initiation of clinical studies.(Kim, Choi et al. 2014) As discussed below. We propose that the splicing modulatory activity may positively contribute to the antitumor activity of all of these clinical drug candidates.

## Results

### An unbiased screen of drugs and drug candidates identifies three investigation antitumor drugs that potently induce the same exon skipping event as the SF3B1 targeted antitumor drug SD6

In order to expand on our initial pilot screen using the TESLR construct integrated into a stable cell-line(Shi, Park et al. 2017) we chose to screen the Selleckchem Bioactive Screening library that is a collection of 2,035 small molecules, which includes FDA approved drugs, compounds that have entered clinical studies and validated tool compounds, see Figure 1. The TESLR construct is stably integrated into an engineered cell line and produces functional luciferase when three exons are skipped in an MDM2 based minigene construct.(Shi, Park et al. 2017) This screen showed a low initial hit rate of 11 hits (see Figure 1) that showed significant activity in replicates (see SI; Table S2), which is consistent with our previous experience with a pilot screen.(Shi, Park et al. 2017) The 11 initial hits were then subjected to dose-response in the TESLR assay (see SI; Figure S1), which confirmed the activity of three compounds (**1**, **2**, and **3**; see Table 1). These three actives were then subjected to confirmation of MDM2 mRNA splicing modulation with RT-PCR using a previously reported gel-based assay that confirmed the expected MDM2 splicing alterations, though the pattern of AS observed with compounds **1**, **2**, and **3** was not identical to that observed with SD6 (see SI Figure S2).(Fan, Lagisetti et al. 2011) The observed splicing changes were more similar between **1**, **2**, and **3** than between that observed with the SF3B1 targeted drug SD6 (see SI Figure S2).

The three confirmed hits were milciclib(Brasca, Amboldi et al. 2009) (compound **1**), PF-3758309(Murray, Guo et al. 2010) (compound **2**) and PF-562271(Roberts, Ung et al. 2008) (compound **3**) (see Table 1). All three of these clinical compounds have published discovery-related selectivity data from different small kinase panels, however none of these panels included members of the CMCG kinase family, which are known to be involved in AS regulation.(Roberts, Ung et al. 2008, Brasca, Amboldi et al. 2009, Murray, Guo et al. 2010). To better understand the breadth of the changes in splicing caused by these compounds we also explored splicing changes in additional mRNAs that are altered by sudemycin D6 using RT-PCR.(Shi, Joyner et al. 2015, Wu, Fan et al. 2018) Our results (See SI; Figure S3) demonstrated that in addition to MDM2 gene, six other genes exhibited clear alternative splicing changes compared to DMSO control, these genes including RBM39, HNRNPR, RAP1B, SRSF7 DUSP, and PAPLOG. We also found that compound **1** and **2** generally have very similar AS pattern to each other, while the splicing pattern changes in some genes when exposed to SD6 or compound **3** were more similar to each other. It should be pointed out that these experiments suggest that the genes which show AS changes within cells exposed to **1**, **2**, or **3** have the potential be used as pharmacodynamic biomarkers for these kinases, as we have previously shown for the SF3B1 targeted agent SD6.(Shi, Joyner et al. 2015, Thurman, van Doorn et al. 2017)

### The three hits are all kinase inhibitors that show potent biochemical and cell-based inhibition of the phosphorylation of substrates

Our previous pilot screening results,(Shi, Park et al. 2017) together with the fact that these three confirmed hits were known to be kinase inhibitors, led us to suspect that these compounds target splicing regulatory kinases. Therefore, we screened these compounds against a panel of kinases known to be involved in the regulation of AS. We then determined the inhibitory IC_50_s for any compound that showed > 50% inhibition at 1 μM for any enzyme in the panel. As shown in Table 1, the confirmed hits showed potent biochemical inhibition of splicing regulatory kinases and other members of the CMCG family, which has not previously been reported for any of these drugs. The relevance of the biochemical data to the cell-based activity of these compounds was demonstrated by measuring total and phosphorylated (phosphor) forms of SR proteins by immunoblots, since these phosphorylation events are known to affect AS (see Figure 2A, 2B). Additionally, we investigated the effect of these drugs on SF3B1 phosphorylation (see Figure 2C, 2D) since a splicing modulatory natural product has been shown to reduce the phosphorylation of SF3B1.(Kumar, Kashyap et al. 2016) 0.5% DMSO was used as control. Interestingly, compounds **1**, and compound **2** reduced phosphorylation of both SR and SF3B1 proteins significantly in HCT116 and Rh18 cell lines, while treatment SD6 did not inhibit the phosphorylation of SR proteins but did inhibit the phosphorylation of SF3B1, which is expected based on the results with pladienolide B in JeKo-1 cells.(Kumar, Kashyap et al. 2016) Compound **3** treatment inhibited the phosphorylation of SR proteins in both HCT116 and Rh18 cells, but to a lesser extent than compound **1** or **2**. Compound **3** also inhibited the phosphorylation of SF3B1 protein in HCT116 cells, but not in Rh18 cells. SR proteins are known to be substrates for CLK and SRPK family members,(Araki, Dairiki et al. 2015) and DYRK1A has been reported to phosphorylate SF3B1(de Graaf, Czajkowska et al. 2006); however, the functional role of SF3B1 phosphorylation is not currently well understood. Based on the kinase inhibition profile of these compounds (Table 1) the reduction of SR protein phosphorylation may be attributed to inhibition of CLKs for compounds **1** and **2**, and the modest inhibition of CLKs by compound **3** may explain the lack of reduction of phosphor-SR protein in cells exposed to this compound. The marked reduction in phosphor SF3B1 by compound **3** points to inhibition of a hypothetical SR kinase that was not included in the panel shown in Table 1. Alternately, it is possible that FAK plays an unrecognized role in the phosphorylation of SF3B1. These results also imply a possible role of SF3B1 phosphorylation in AS, that can be the subject of further experiments.

**Figure 1.**
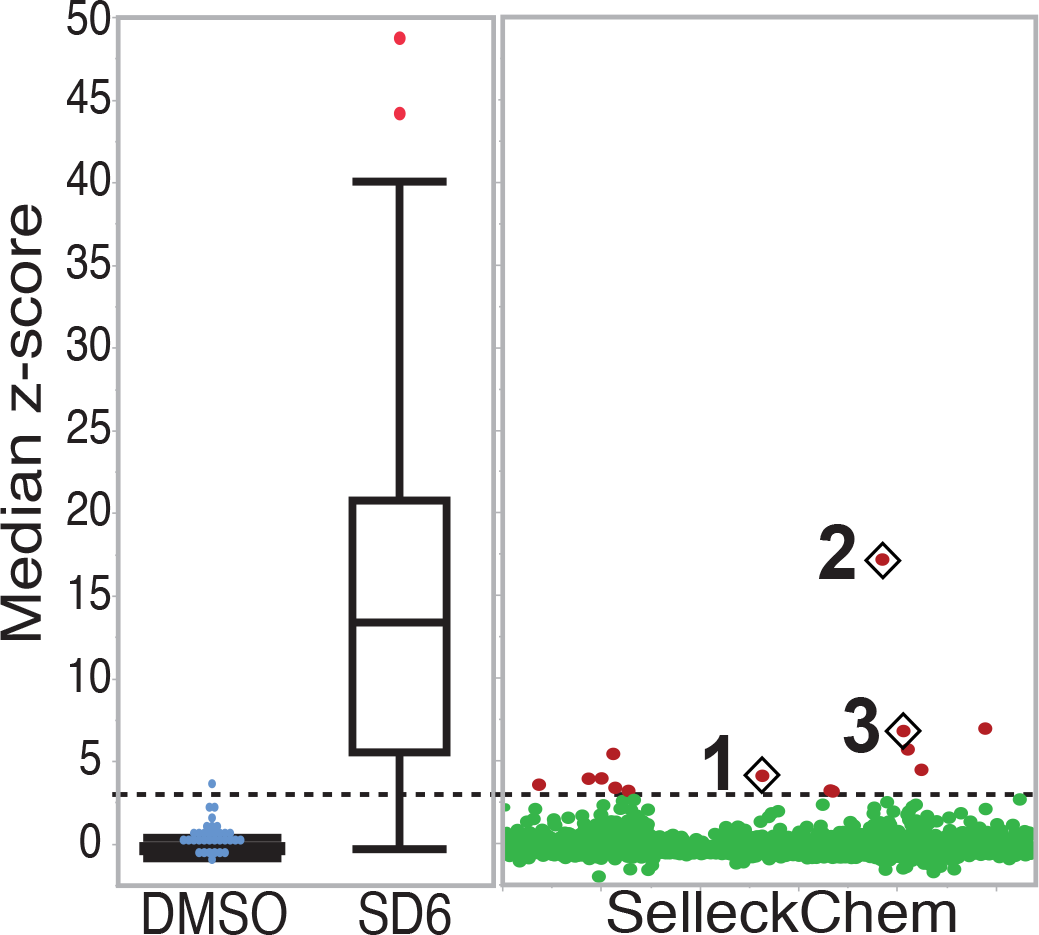
Screening results showing median Z-scores from three replicates, with the combined negative (DMSO) and positive (SD6) controls shown on the left, and the complete dataset from the SelleckChem library on the right. Shown in red are the compounds that were selected for follow-up dose-response studies based on their median Z-scores (>3), and in diamonds are the three compounds that showexsd a dose-response upon retesting.

**Table 1.**
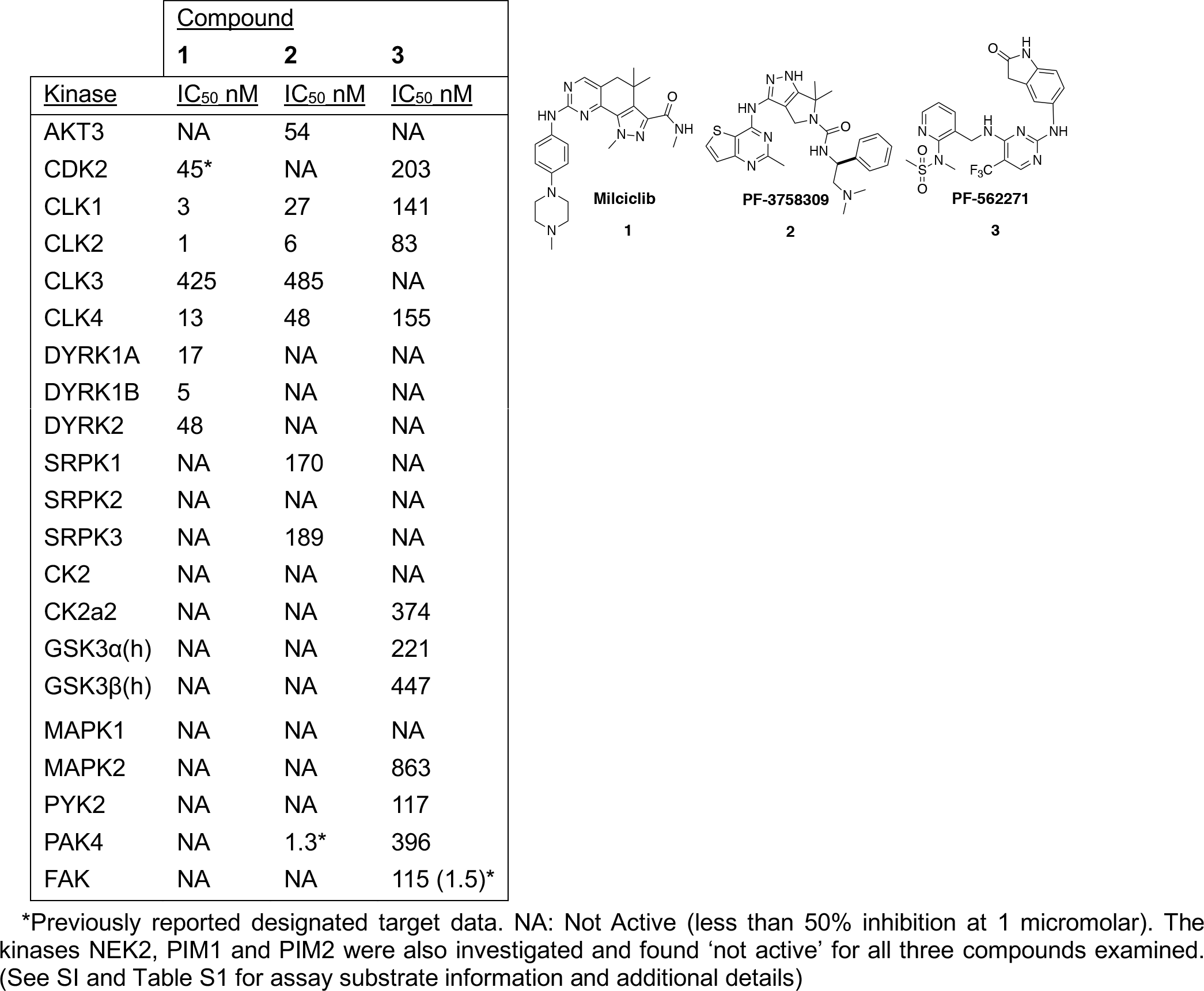
Biochemical kinase inhibition data of the confirmed TESLR hits with structures

**Figure 2.**
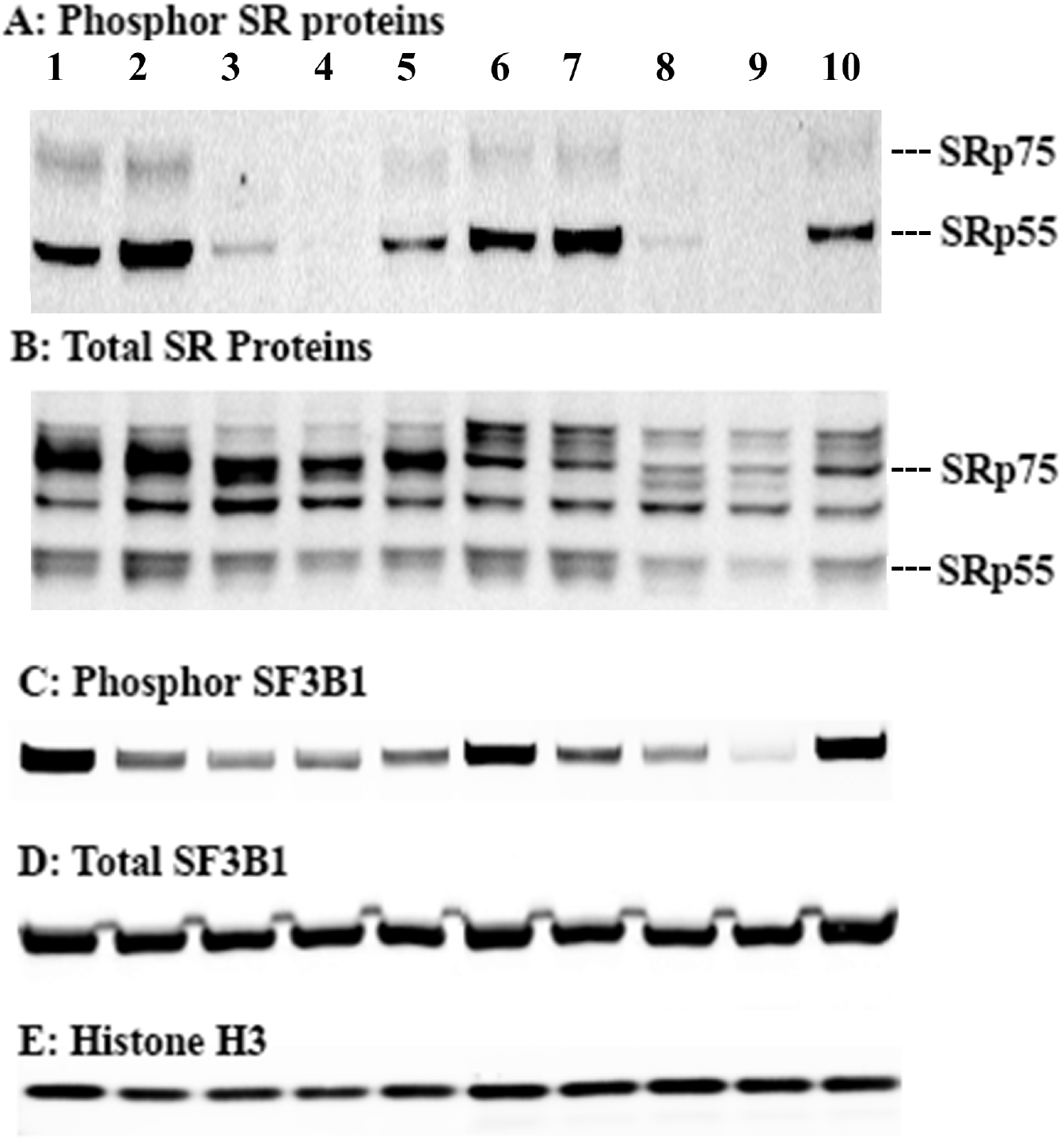
Nuclear extracts blotted with antibodies to phosphor-SR, total SR, phosphor-SF3B1, total SF3B1 or histone H3 proteins from HCT-116 cells and Rh18 cells treated with DMSO, SD6 or compounds **1**, **2**, or **3** for 4 h. Lanes for all gels: Left to right: Lanes 1 to 5 are nuclear extracts from HCT116 cells. Lane 1: DMSO; Lane 2: 10 μM SD6; Lane 3: 10 μM compound **1**; Lane 4: 5 μM compound **2**; Lane 5: 10 μM compound **3**; Lanes 6 to 10 are nuclear extracts from Rh18 cells. Lane 6: DMSO, 4 h; Lane 7: 10 μM SD6; Lane 8: 10 μM compound **1**; Lane 9: 5 μM compound **2**; Lane 10: 10 μM compound **3**. These blots are representative of three independent experiments. Procedures can be found in the SI in supporting section G.

### Focused screening of selective tool compounds and kinase inhibitory drugs uncovered additional active compounds in the TESLR screen that inhibit a subset of splicing regulatory kinases DYRK, CLK, or CK but not SRPK 1 or 2

Given the potent and previously unrecognized splicing kinase inhibitory activity of these drugs against this kinase panel, we also decided to investigate the activity of other known splicing kinase inhibitory drugs and tool compounds that were not included in the initial TESLR screen. To this end we performed a dose-response screen with Mirk-IN-1 (a known DYRK1A inhibitor),(Anderson, Chen et al. 2013) Sphinx31 (a selective SRPK1 inhibitor)(Batson, Toop et al. 2017) dinaciclib (a FDA designated orphan drug and a potent CDK 1, 2, 5, and 9 inhibitor),(Paruch, Dwyer et al. 2010) SRPIN340 (SRPK 1 and 2 inhibitor),(Karakama, Sakamoto et al. 2010) SNS-032 (CDK 2,7 and 9 inhibitor),(Ma and Cress 2007) flavopiridol (Alvocidib, a FDA designated orphan drug and a potent pan-CDK inhibitor),(Senderowicz 1999) palbociclib (a non-selective CDK/multi-kinase inhibitor that is a FDA designated orphan drug),(Klaeger, Heinzlmeir et al. 2017) and silmitasertib (a CK2 inhibitor in Phase II).(Kim, Choi et al. 2014) Of these compounds we found that only Mirk-IN-1, palbociclib and silmitasertib were active (IC_50_s in the range of 10-300 μM) in the TESLR assay, while the other splicing kinase inhibitors showed IC_50_s >> 10 μM in inducing triple exon skipping (see SI; Figure S4). This shows that the type of alternate splicing detected by the TESLR assay is not induced by SPRK inhibitors and also indicates that the inhibition of the DYRKs, casein kinase 2 (CK2), or CLKs 1, 2, and/or 4 (and the resulting inhibition of phosphorylation of SR proteins or SF3B1) were the likely causes of the triple-exon skipping observed in the TESLR screen with compounds **1**, **2**, **3**, and palbociclib, although the possibility that inhibition of FAK or other non-designated splicing kinases may also be involved, was not excluded by these studies. These results are completely consistent with recent findings from the chemoproteomic analysis of 243 kinase inhibitory drugs and tool compounds, which showed that drugs (e.g. palbociclib) and tool compounds that were touted as ‘selective’ actually potently inhibit numerous kinases, in addition to their designated targets.(Klaeger, Heinzlmeir et al. 2017) Notably this comprehensive work clearly shows that palbociclib (trade name: Ibrance) has some affinity for CLK1 (IC_50_ = 276 nM),(Klaeger, Heinzlmeir et al. 2017) which presumably accounts for the modest splicing modulation observed with this approved antitumor drug.

### Interrogating the SF3B1 small molecule binding site: the development of a competitive antagonist assay

In order to better understand the activity profile of TESLR hits we also decided to develop an improved assay for SF3B1 small molecule binding, since it is always possible that these hits could directly interact with SF3B1. We therefore decided to take advantage of the novel observation from the Jurica lab that ‘inactive’ analogs of natural product SF3B1 ligands can displace potent natural product splicing modulators in cell-free *in vitro* systems.(Effenberger, Urabe et al. 2016) Since this report indicated that it was possible that our previously synthesized natural product analogs could be SF3B1 antagonists we screened a set of 14 of ‘inactive’ (minimal to no TESLR activity and non-cytotoxic at the concentrations investigated) sudemycin and herboxidiene analogs in the TESLR assay in the presence of moderate concentrations of SD6 (see Figure 3; and SI, Tables S4 and S5). This assay identified two compounds that were able to potently antagonize the activity of SD6 as shown in Figure 3. Using this assay format, we were then able to show that sudemycinol C did not antagonize the TESLR activity of compounds **1**, **2**, or **3**, which indicates that these hits do not bind to the same site on SF3B1 as SD6 (see SI; Figure S3), which is consistent with our proposal that the inhibition of subsets of splicing regulatory kinases accounts for the splicing modulatory activity of these drugs. These results represent the first demonstration of SF3B1 antagonism in cells and the first HTS assay useful for the determination of binding to the splicing modulator binding site of SF3B1.

**Figure 3.**
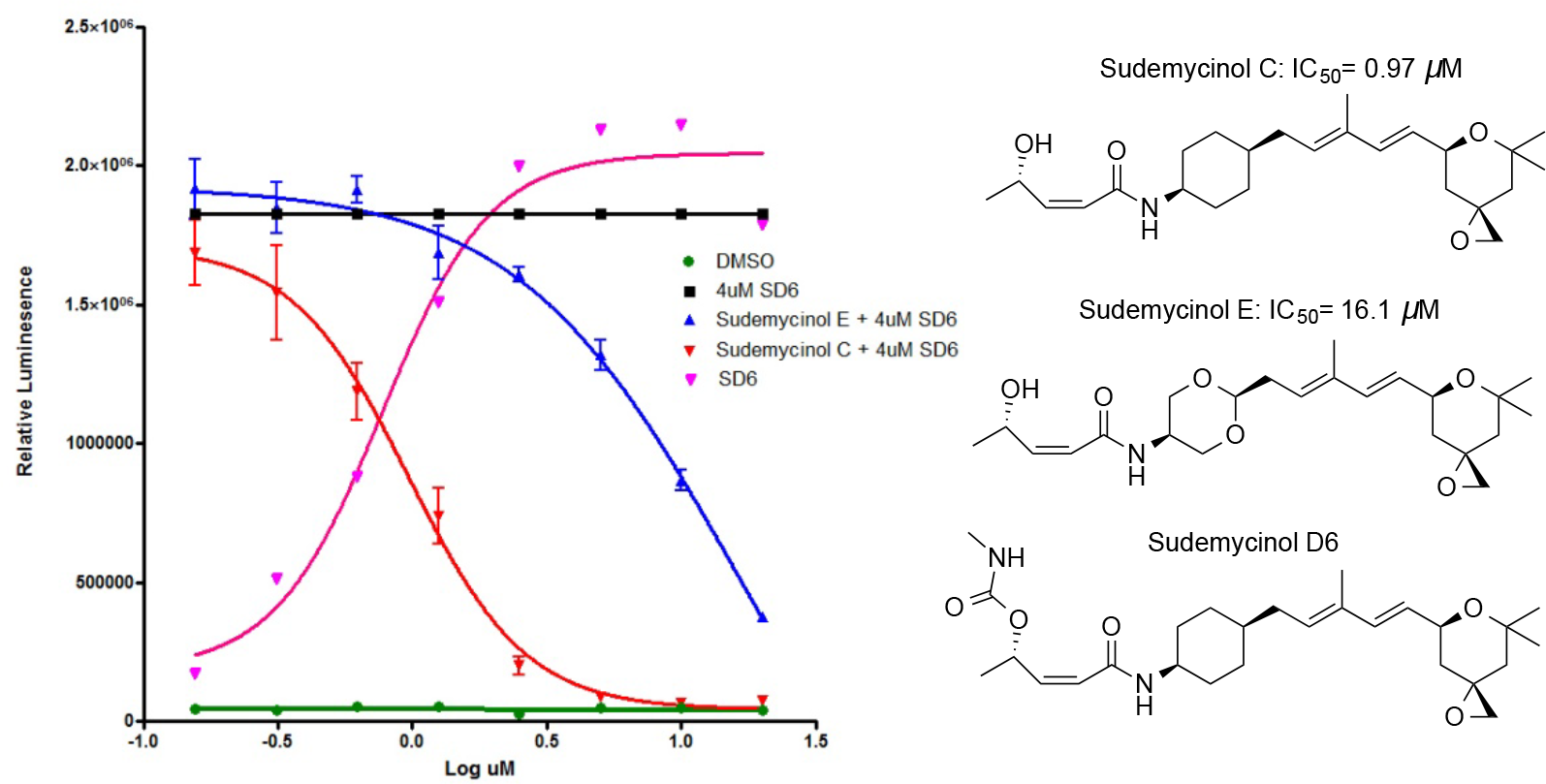
(Left Panel) The assay that identified two antagonists of SD6 activity in the TESLR screen, which are useful for the determination of SF3B1 binding. Relative luminescent units were plotted against corresponding drug concentrations and fitted with a standard four parameter sigmoidal curve with GraphPad Prism. (Right Panel) The structures of the antagonists and sudemycin D6 (SD6). See SI section F for details.

## Discussion

One major goal of this work was the discovery of new splicing modulatory drug-like lead compounds that exhibit similar splicing changes to those observed with the SF3B1 targeted natural products and analogs. To accomplish this, we developed the TESLR screen that was designed based on a prevalent, unusual and characteristic modification of pre-mRNA splicing we observed in tumor cells treated with SF3B1 antitumor natural products and analogs. We are therefore pleased to identify three post-Phase I antitumor drugs that may exhibit a substantial component of their activity through the modulation of splicing, in combination with their designated target engagement. It is interesting to note that though all available drugs and available clinical compounds were included in the screen, only antitumor drugs were identified, though this could simply be due to the fact that large number of kinase inhibitors are under clinical investigation and many investigational antitumor drugs are kinase inhibitors that typically exhibit polypharmacology.(Klaeger, Heinzlmeir et al. 2017) Naturally, these results suggest that the repurposing of these multitargeted drugs should be envisioned in light of this new clinically-relevant MOA information. These results also clearly highlight the need for careful scholarship and scrutiny when using ‘selective’ drugs as tool compounds for experiments in cell biology.(Arrowsmith, Audia et al. 2015) Fortunately, an authoritative publication of inhibitor binding data for 243 clinical kinase inhibitors with 220 human kinases makes this important kinase selectivity data readily available to the biomedical science community.(Klaeger, Heinzlmeir et al. 2017)

In order to evaluate the possible direct interaction of **1**, **2**, or **3** with the small molecule binding site of the SF3B subunit, we screened a small focused library of analogs of FR-901,464 and herboxidiene that show minimal activity in the TESLR assay, at the concentrations examined. Two of these compounds (sudemycinol C and E) were able to effectively reduce the TESLR activity of SD6 in a dose-dependent manner, presumably by competing with SD6 for the SF3B1 binding site without triggering the modulation of splicing detected by the TESLR assay. This observation is consistent with results from cell-free experiments, reported from the Jurica laboratory, which showed that binding to the small molecule SF3B1 site is necessary, but not sufficient, to induce splicing modulation and reported several different antagonists at this site.(Effenberger, Urabe et al. 2016) Thus we discovered new simple antagonists and extended this type of assay into a format using living cells, which is useful for the high-throughput determination of SF3B1 small molecule binding. Using this screen, we were able to show that the activity seen with **1**, **2**, or **3** is not affected by the antagonist sudemycinol C, which is consistent with our hypothesis that these drugs exert their action through the inhibition of splicing modulatory kinases.

In summary, we report the novel characterization of three drugs as splicing modulators **1** (milciclib), **2** (PF-3758309), and **3** (PF-562271) that show potency on par with the SF3B1 targeted drug candidate sudemycin D6 in the TESLR screen, which reports on a triple-exon skipping event in MDM2 pre-mRNA that we first observed in tumor cells treated with sudemycin analogs.(Fan, Lagisetti et al. 2011) We further demonstrated that in addition to modulation of splicing of the MDM2 gene these three drugs differentially induce AS in multiple genes that we examined, further supporting our hypothesis that the splicing modulatory activity is an important aspect of the antitumor activity of these three drugs. We also characterized the MOA of these drugs as the potent biochemical inhibition of a subset of the CMCG family of kinases including DYRK, CLK and CK. Additionally, we confirmed that these compounds show cell-based inhibition of the phosphorylation of SR proteins and SF3B1. Though the functional role of phosphorylation of SF3B1 is not currently fully understood,(de Graaf, Czajkowska et al. 2006) our results suggest a possible regulatory role of this phosphorylation event on alternative splicing, and show that the SF3B1 protein appears to be a substrate for other kinases including the CLKs.

We also report the discovery of new potent SF3B1 antagonists (sudemycinol C and sudemycinol E) for the SF3B1-sudemycin binding site, and thereby confirm the previous report of the remarkable pharmacology of this critical spliceosome regulatory component.(Effenberger, Urabe et al. 2016) We also show that these newly identified antagonists allow for a facile determination of the compounds that interact with the SF3B1 SD6 binding site, via a simple competition assay coupled to the TESLR splicing reporter screen. Using this new assay, we show that hit compound’s splicing modulatory activity in TESLR is not mediated through binding to the SF3B1 small molecule binding site, which is consistent with our hypothesis that they act through their potent inhibition of different splicing factor kinases. Our results also highlight a previously unanticipated portrait of the receptor-like pharmacology of SF3B1 splicing protein that is beginning to emerge.

These new insights suggest novel opportunities for the future clinical repositioning of these agents for additional indications in oncology, since these drugs have already been evaluated through Phase I or Phase II clinical studies. This improved understanding of the MOA of these clinical agents can lead to the effective translation of basic research results into better informed future clinical studies with these drugs, with an increased likelihood of therapeutic success. Specifically, our work suggests that the pre-clinical efficacy of compounds **1**, **2**, and **3** should be examined for possible clinical repurposing in MDS, CLL and AML. Our work also adds new tools compounds, and further clarifies the activity of known splicing modulatory compounds for basic research in cell biology. These interconnected outcomes support a growing recognition of the substantial potential of modulators of pre-mRNA splicing in cancer therapy, and also suggest a profile for a new chemotherapeutic class of multi-targeted splicing kinase inhibitors with other potential clinical applications, which is of interest to scientists across a broad range of disciplines from drug discovery through clinical practice in oncology.

## Materials and Methods

High Throughput Screening Methods: SK-Mel-2-MDM2-Luc cells were cultured in MEM supplemented with 10% fetal bovine serum, 1 mM pyruvate, and 0.1 mg/mL of the antibiotic G418; the cells were grown in a 37°C incubator at 5% CO_2_.(Shi, Park et al. 2017) To perform the TESLR high-throughput screen, white 384 plates were seeded with 20 µL per well of growth medium without G418, at a density of 5,000 cells per well using a Multidrop peristaltic pump (Thermo Scientific). After 18 h at 37°C in an incubator in 5% CO_2_, cells were treated with 100 nL of compound using a pin tool attachment for a Janus MDT automated workstation (Perkin Elmer). SD6 at 5 µM was used as positive control for splicing modulation. After 5 h incubation at 37°C with 5% CO2, an equal volume of OneGlow XL (Promega) was added using a Matrix WellMate dispenser (Thermo Scientific). Luminescence readings were taken immediately using an Envision multilabel reader (PerkinElmer). We screened 2035 compounds (seven 384-well plates) from SelleckChem at 10 µM in triplicate. The first 2 columns were reserved for the SD6 positive control, while the last two columns contained DMSO as a negative control. Z-scores were calculated from the DMSO-treated negative controls using the following equation: [X – µ(DMSO)]/σ(DMSO), where X = the luminescence intensity for a given well, and µ(DMSO) and σ(DMSO) are the average and standard deviation of the DMSO (calculated for each plate separately). Hits were determined as those wells having a median Z-score greater than 3. There were 12 hits that scored as positives by these criteria in both replicates, and these were repurchased, their purities confirmed by LCMS, and submitted to dose-response testing for confirmation of splicing modulation activity and to determine potency. All other experimental details are included in the Supporting Information section.

## ASSOCIATED CONTENT

### SUPPORTING INFORMATION

General methods for synthetic chemistry and biochemical assays (including kinase substrate information), list of initial hits, dose response curves and RT-PCR gels for hit confirmation and tool compound evaluation, details on antagonist screen and binding assay results with hits. Experimental details for the synthesis of the unpublished screened natural product analogs.

## Author Contributions

Y.S. assisted with the HTS, performed all hit confirmation dose response curves and developed the antagonist screen and assay. W.B. and A.S. performed the HTS screen and initial hit identification, which was supervised by S.L. J.C. ran and processed the Western blots. C.L. provided the natural product analogs for the antagonist screen. W.Z. was responsible for the kinase activity profiling. L.S. led the cell biology, interpreted the Western blots and prepared Figure 2. T.R.W. led most of the chemical biology experimental design. All authors contributed to the writing or editing of the manuscript and have given approval to the final version of the manuscript.

## Notes

The authors declare no competing financial interests.

## ACKNOWLEDGMENTS

The authors wish to thank Karen Tenney of the Department of Chemistry and Biochemistry at UCSC for her contributions to the administration and organization of this project. We also gratefully thank Professor Melissa Jurica of the Department of Molecular, Cell & Developmental Biology at UCSC, for her very helpful critical review and edits to the draft manuscript.

## Abbreviations

MOA: mechanism of action
IND: investigational new drug
TESLR: triple exon skipping luciferase reporter
SD6: sudemycin D6
HTS: high-throughput screening
AS: alternative splicing
CLL: chronic lymphocytic leukemia
MDS: the myelodysplastic syndromes
Phosphor: phosphorylated

