## Supplementary Information for "An Exon Skipping Screen Identifies Antitumor Drugs That Are Potent Modulators of Pre-mRNA Splicing, Suggesting New Therapeutic Applications"

---

### SUPPORTING INFORMATION

#### Table of Contents

**A. General Experimental Methods** **Page 2-3**

**B. List of Initial Hits** **Page 4**

**C. TESLR Dose Response Hit Confirmation** **Page 4-6**

**D. Confirmation of Splicing Changes With RT-PCR** **Page 6-9**

**E. TESLR Dose Response Tool Compounds** **Page 9-10**

**F. Antagonist Screen and Assay for Confirmed Hits** **Page 10-11**

**G. Immunoblotting Analysis** **Page 11-12**

**H. Natural Product Analog Synthetic Chemistry** **Page 12-15**

### A. General Methods

#### Biochemical kinase inhibition assays.

Enzymatic biochemical activities were evaluated in radiometric protein kinase assays (Eurofins, Dundee, Scotland). Kinases are incubated with buffer, substrate and [γ-$^{33}$P]-ATP (specific activity approx. 500 cpm/pmol, concentration as required). The reaction is initiated by the addition of the MgATP mix. After incubation for 40 minutes at room temperature, the reaction is stopped by the addition of 3% phosphoric acid solution. 10 µL of the reaction is then spotted onto a P30 filtermat and washed three times for 5 minutes in 75 mM phosphoric acid and once in methanol prior to drying and scintillation counting. All compounds are prepared to 50x final assay concentration in 100% DMSO. Positive control wells contain all components of the reaction with 2% DMSO. Blank wells contain all components of the reaction, with a reference inhibitor replacing the compound of interest. Each kinase is assigned a standard assay concentration of ATP within 20 µM of its apparent $K_m$. Compounds were screened at 1 µM in duplicate or tested at 10 concentrations at half log dilution starting from 10 µM in singlicate. Dose response curves were fitted with four parameter logistic curve to obtain $IC_{50}$ values. The average coefficient of variation (CV) of the conducted assay.

Percent inhibition was calculated by comparing to the positive control wells that contain all components of the reaction and 2% DMSO instead of compound (0% inhibition), as well as the blank wells that contain all components of the reaction, with a reference inhibitor (100% inhibition). Staurosporine is used as reference for all tested enzymes except for SRPK3 (reference inhibitor: K-252a. Nocardiopsis sp), NEK2 (reference inhibitor: 30% phosphoric acid), and CK2, CK2α2, MAPK1, MAPK2 (reference inhibitor: PKR Inhibitor). See Table S1 below for additional details.

| Enzyme[a] | Substrate (concentration) | ATP (µM) | %CV[g] |
|---|---|---|---|
| AKT3 | GRPRTSSFAEGKK (30µM) | 45 | 3 |
| CDK2/cyclinA (h) | histone H1 (0.1 mg/mL) | 45 | 6 |
| CLK1(h)[b] | ERMRPRKRQGSVRRRV (200µM) | 10 | 2 |
| CLK2(h) | YRRAAVPPSPSLSRHSSPHQS(p) EDEEE (20µM) | 10 | 4 |
| CLK3(h)[c] | ERMRPRKRQGSVRRRV (250µM) | 90 | 4 |
| CLK4(h) | YRRAAVPPSPSLSRHSSPHQS(p)EDEEE (200µM) | 10 | 3 |
| DYRK1A(h) | RRRFRPASPLRGPPK (50µM) | 45 | 7 |
| DYRK1B (h) | RRRFRPASPLRGPPK (50µM) | 45 | 4 |
| DYRK2 (h) | casein (2mg/ml) | 15 | 3 |
| SRPK1(h) | RSRSRSRSRSRSRSR (300µM) | 10 | 10 |
| SRPK2 (h) | RSRSRSRSRSRSRSR (300µM) | 10 | 4 |
| SRPK3 (h) | ERMRPRKRQGSVRRRV (50µM) | 15 | 7 |
| CK2 (h)[d] | RRRDDDSDDD (165µM) | 15 | 5 |
| CK2α2 (h)[d] | RRRDDDSDDD (330µM) | 10 | 3 |
| GSK3α(h) | YRRAAVPPSPSLSRHSSPHQS(p) EDEEE (20µM) | 10 | 6 |
| GSK3β(h) | YRRAAVPPSPSLSRHSSPHQS(p) EDEEE (20µM) | 15 | 6 |
| MAPK2 (h)[e] | myelin basic protein (0.33 mg/mL) | 155 | 5 |
| PYK2 (h) | poly(Glu, Tyr) 4:1 (0.1 mg/mL) | 90 | 5 |
| PAK4 (h) | myelin basic protein (0.8 mg/mL) | 10 | 7 |
| FAK (h)[f] | EEEEYEEEEEEYYIIEEEEEEYEEEEEEYYEEEEEEKKKK (100µM) | 70 | 5 |
| NEK2 (h) | myelin basic protein (0.33 mg/mL) | 120 | 4 |
| Pim-1 (h) | KKRNRTLTV (100µM) | 90 | 4 |
| Pim-2 (h) | RSRHSSYPAGT (300µM) | 15 | 10 |

**Table S1.** Substrates and conditions for biochemical $IC_{50}$ determinations.

[a] The standard enzyme reaction buffer contains mM MOPS pH 7.0, 10 mM Mg Acetate, 0.2 mM EDTA. The additional components of any specific enzyme are described from b-k.
[b] The enzyme reaction buffer for CLK1(h) also contains 1 mM sodium orthovanadate, 5 mM sodium 6-glycerophosphate.
[c] The enzyme reaction buffer for CLK3(h) also contains 30 mM NaCl.
[d] The enzyme reaction buffer for CK2 (h) and CK2α2 (h) also contain 20 mM HEPES, 0.15 M NaCl, 0.1 M EDTA, 5 mM DTT, 0.1% Triton X-100, 50% Glycerol.
[e] The enzyme reaction buffer for MAPK1 (h) and MAPK2 (h) also contain 50 mM TRIS, 0.1 mM EGTA, 0.1 mM $Na_3VO_4$, 0.1% 6-mercaptoethanol, 1 mg/mL BSA.
[f] The enzyme reaction buffer for FAK (h) also contains 15 mM MOPS, 0.75 mM EDTA, 0.0075% Brij-35, 3.75% Glycerol, 150 mM NaCl, 0.1% 6-mercaptoethanol, 1 mg/mL BSA.
[g] Percent CV is of the assay is calculated from the CV of four control compound repeats across different experiments.

### Chemistry

*General.* Unless otherwise noted, all commercial reagents were obtained from commercially available sources and used without purification. Flash column chromatography was performed on a Biotage SP-1 chromatography system. TLC plates were visualized by exposure to ultraviolet light (254 nm). $^{1}H$ and $^{13}C$ spectra were recorded using 400 MHz, and 300 MHz, respectively, using $CDCl_3$, $CD_3OD$, or DMSO-$d_6$ as a solvent. The chemical shifts are reported in parts per million (ppm) relative to residual solvent (for chloroform, $\delta$ 7.24 ppm for $^{1}H$ NMR and $\delta$ 77.02 ppm for $^{13}C$ NMR. For DMSO, $\delta$ 2.47 ppm for $^{1}H$ NMR. For $CD_3OD$, $\delta$ 49.00 ppm for $^{13}C$ NMR). Coupling constants are reported in hertz (Hz). The following abbreviations are used to designate the multiplicities: s = singlet, d = doublet, t = triplet, q = quartet, m = multiplet. Mass spectra with electrospray ionization (ESI) were recorded on LCQ Fleet Ion Trap Mass Spectrometer (Thermo Scientific) coupled to the Finnigan Surveyor Plus HPLC System (Thermo Scientific). High-resolution mass spectra were recorded on LTQ-Orbitrap XL (Thermo Scientific) using static nanoelectrospray ionization in positive-ion profile mode at a nominal resolution setting of 100,000. Approximately 50 scans were averaged for each sample and the resulting Fourier-transformed frequency-domain spectrum was mass-assigned with calibration constants from an external calibration mixture. Experimental masses and isotope distributions were compared to theoretical values. All compounds reported are of at least 95% purity, as judged by HPLC (Waters XBridge C18, 250 mm X 4.6 mm ID, 5 µm column; 10 µL injection; 10-100% MeCN/$H_2O$ + 0.1% TFA gradient over 15 min; 1 mL/min flow; ESI; positive ion mode; UV detection at a wavelength of 310 or 340 nm).

### B. List of Initial Hits

**Table S2**. The initial hit compounds; SRI ID, compound name, purity and molecular weight. These compounds are all commercially available. See Section C below for TELSR dose response curves.

| SRI Number | Compound Name | Batch Purity | MW |
|---|---|---|---|
| SRI-030859 | Urapidil | >95% | 387 |
| SRI-030863 | Chlorothiazide | >95% | 295 |
| SRI-030864 | Olmesartan | >95% | 558 |
| SRI-030865 | Bekanamycin | >95% | 483 |
| SRI-030866 | Birinapant | >95% | 806 |
| SRI-030867 | Milciclib | >95% | 460 |
| SRI-030868 | PF-3758309 | >95% | 490 |
| SRI-030869 | Ansamitocin P-3 | 76% | 635 |
| SRI-030870 | AZD6482 | >95% | 408 |
| SRI-030871 | PF-562271 | >95% | 665 |
| SRI-030872 | Clevidipine Butyrate | >95% | 456 |

### C. TESLR Dose Response

*Experimental*: SK-MEL-2/MDM2-Luc stable cells were cultured in MEM medium with Earle's salts and L-glutamine containing 1 mM sodium pyruvate, 10% FCS and 10 mM Hepes and plated at a density of 10,000/well in 96-well plates and incubated overnight at 37°C in 5% $CO_2$. The following day, cells were treated with serial dilutions of compounds for 4 hrs, ONE-Glo reagents (Promega) were added to measure the luciferase activity on an EnVision plate reader. Sudemycin D6 and 0.5% DMSO were used as positive and negative controls, respectively. Relative luminescent units were plotted against corresponding drug concentrations and fitted with a standard four parameter sigmoidal curve with GraphPad Prism.

### Hit Confirmation

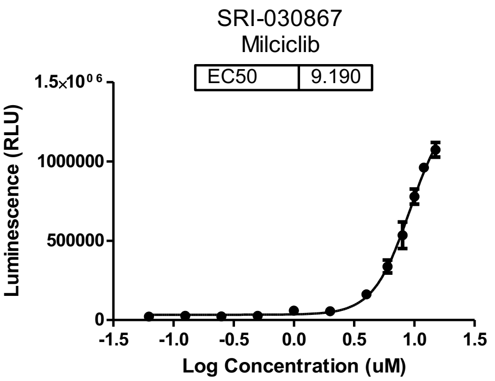


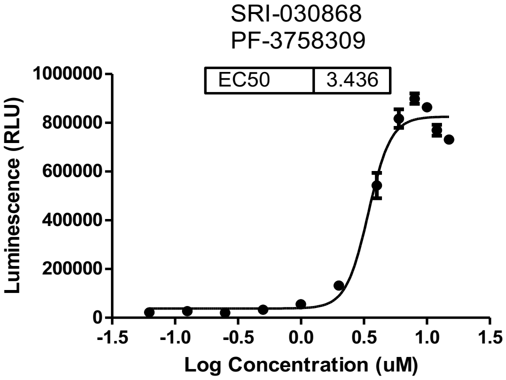


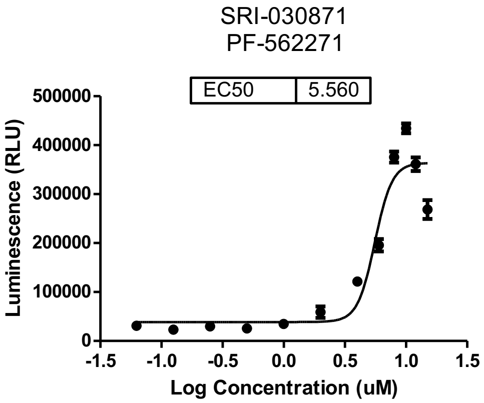


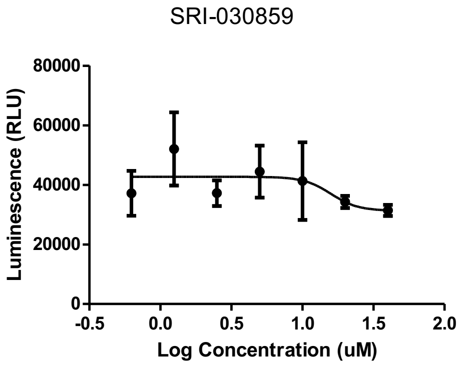


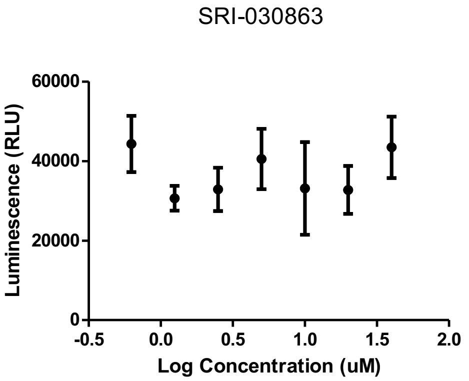


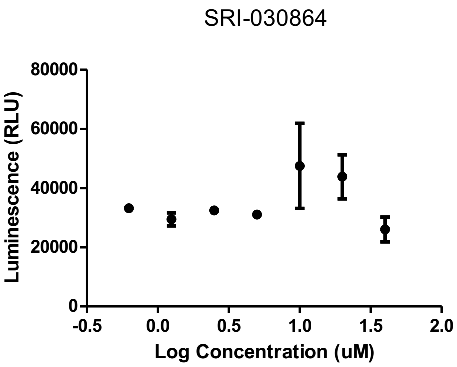


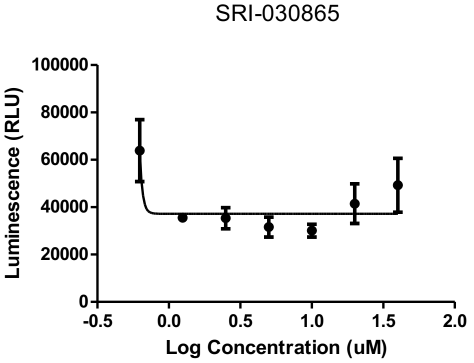


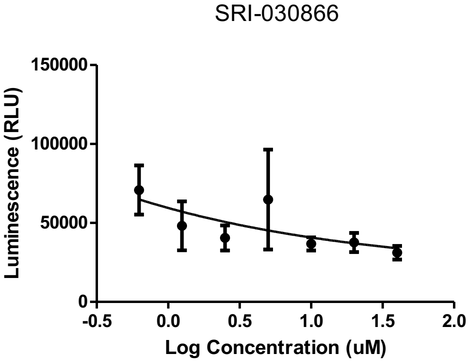

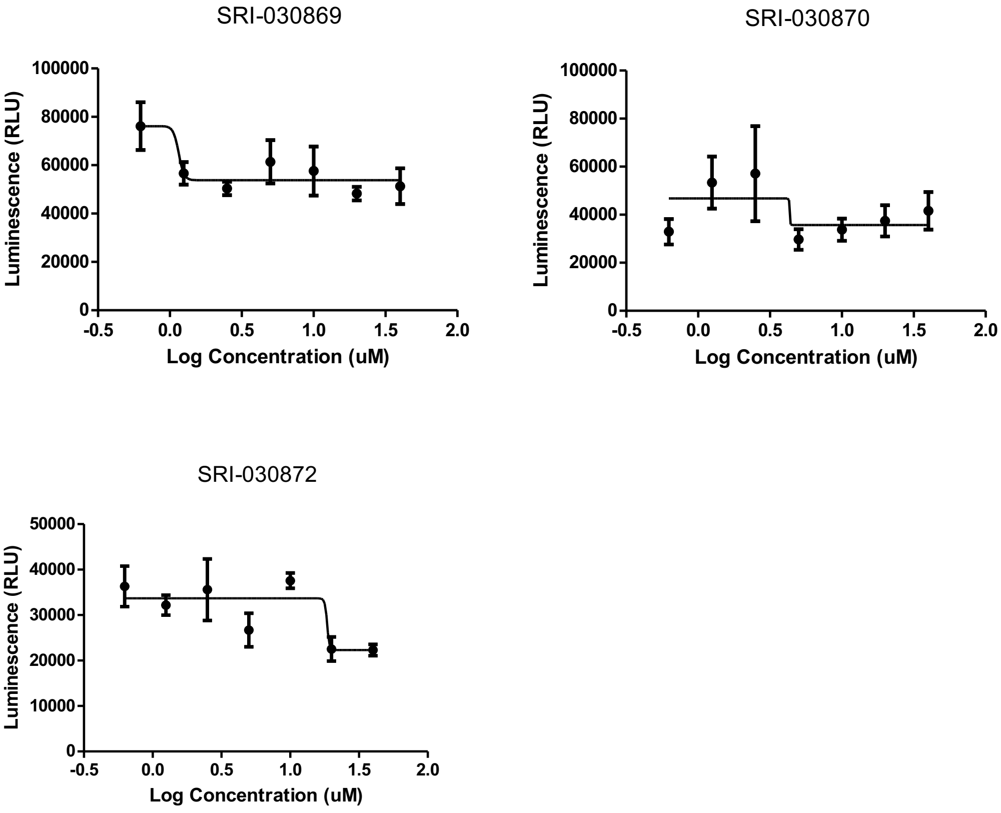


**Figure S1.** Dose response curves of all initial hits identified from the library screen. TESLR assay in SK-MEL-2/Luc-MDM2 cells with 4 hours treatment. SK-MEL-2/Luc-MDM2 stable cells were plated at a density of 20,000 cells/well in 96-well plates and incubated overnight at 37°C in 5% $CO_2$. The following day, cells were treated with either 0.5% DMSO or serial dilutions of hits identified from the library screen in 0.5% DMSO for 4 hours. ONE-Glo™ EX reagent (Promega) was added to measure the luciferase activity using an EnVision plate reader. Relative luminescent units were plotted against corresponding drug concentrations and fitted with a standard four parameter sigmoidal curve with GraphPad Prism. $EC_{50}$ values are shown for the three compounds that produced a dose response curve.

### D. Confirmation of Splicing Changes with RT-PCR

*Experimental*: RH-18 cells were cultured in RPMI 1640 medium containing 10% FCS. Cells are all maintained in humidified incubator with 5% CO2 at 37°C. Rh18 cells were exposed to 0.5% DMSO or sudemycin D6 (SD6) 10 µM , Cpd1 10 µM, Cpd2 5 µM or Cpd3 10 µM for 4 h. Total RNA was extracted and converted to cDNA. PCR was performed using NEB Q5 Master Mix and specific primers listed below (Table S3) according to the manufacturer's instruction and standard PCR protocols: 50 ng cDNA and 30 µl of the final reaction volume was used. PCR products were subjected to 3% agarose gel electrophoresis.

**Table S3**. List of primers used in the RT-PCR.

| primer | sequence |
|---|---|
| MDM2 Forward | CTGGGGAGTCTTGAGGGACC |
| MDM2 Reverse | CAGGTTGTCTAAATTCCTAG |

| | |
|---|---|
| UBQ Forward | ACCTGACCAGCAGCGTCTGATATT |
| UBQ Reserve | TCGCAGTTGTATTTCTGGGCAAGC |
| DUSP 11 Forward | GACATCAAGTGCCTGATGATGA |
| DUSP11 Reverse | ATGTCCCCGGCACCTATT |
| SRRM1 Forward | GACTCTGGCTCCTCCTCCTC |
| SRRM1 Reverse | GGACTTCTCCTCCGTCTACCA |
| MLH3 Forward | TTATTGCCTGTTTGATGAGCAC |
| MLH3 Reverse | TCCTTTGTTCCTCTGTCACTGTT |
| PAPOLG Forward | AAGAGATCCCATTCCCCATC |
| PAPOLG Reverse | TGCGTGATGTATCAATAGTTGGA |
| HERPUD1 Forward | GTCGTTGCAGAGATTGCGGG |
| HERPUD1 Reverse | GCATATATCTGCTGGAACCAGG |
| HNRNPR Forward | GAGCAATTGATGCTCTCAGG |
| HNRNPR Reverse | TCTGTCACCGTAGTAGCCAAG |
| RAC1 Forward | ATGCAGGCCATCAAGTGTGTG |
| RAC1 Reverse | GTTCCAAGGGACAGGACCAAG |
| RAN Forward | TCCGCCATCTTTCCAGCCTCA |
| RAN Reverse | GAATTTCTCCTGGCCGGCTG |
| RAP1B Forward | AGGAGGCGTTGGAAAGTCTG |
| RAP1B Reverse | GGCACTGTTGAATTGGGCAAC |
| RBM39 Forward | GAAGCAATGCTTGAGGCTCC |
| RBM39 Reverse | GCTCAAAGATCCCACGAAGC |
| SMEK1 Forward | GGATCCCGCTACAATCTGATG |
| SMEK1 Reverse | CTACAGCTTCTCCATCTTCC |
| SRSF7 Forward | GAACTATCGACAGGCATGCC |
| SRSF7 Reverse | CTTCTGCGAGGACTTCCTGAT |
| VDAC3 Forward | GACTTCCAGCTGCACACACA |
| VDAC3 Reverse | CCAGGGACAAATCTGATGC |

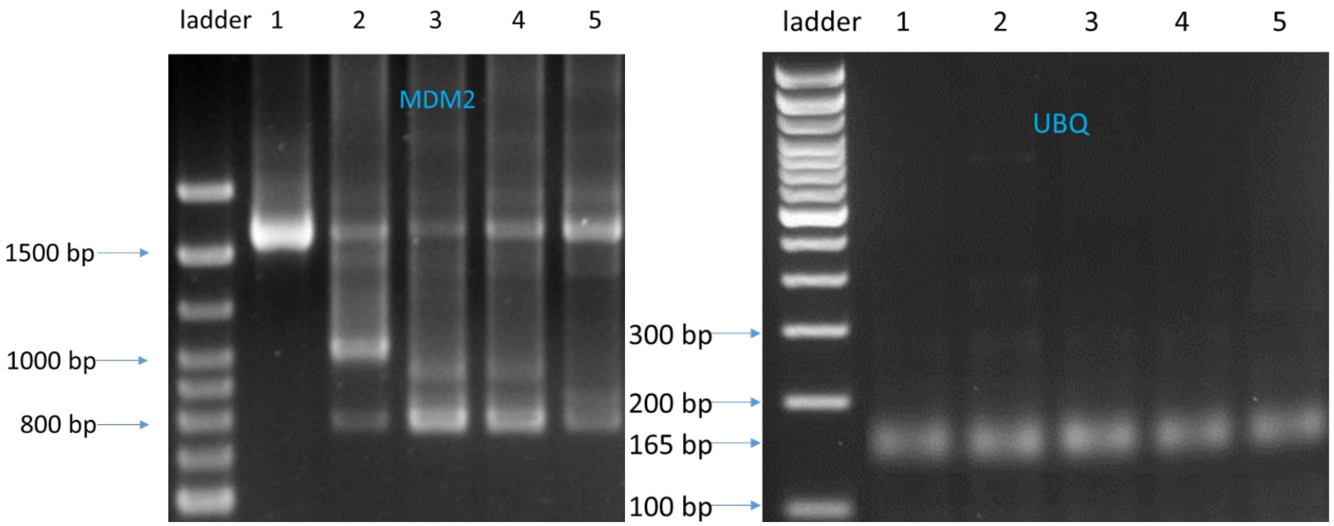


Figure S2. Effects of SD6, **1**, **2**, and **3** on pre-mRNA splicing for the MDM2 gene in Rh18 using RT-PCR. (Lane 1) DNA Ladder; (Lane 2) 0.5% DMSO; (Lane 3) SD6: 10 µM; (Lane 4) **1**: 10 µM; (Lane 5) **2:** 5 µM; (Lane 6) **3**: 10 µM. Rh18 cells were treated for 4 h. Total RNA was extracted and converted to cDNA.

PCR was performed by using corresponding primers and PCR products were subjected to 3% agarose gel electrophoresis. AS was identified with compound treatments for MDM2 transcripts (Left). Ubiquitin (UBQ) transcripts were used as a control (Right).

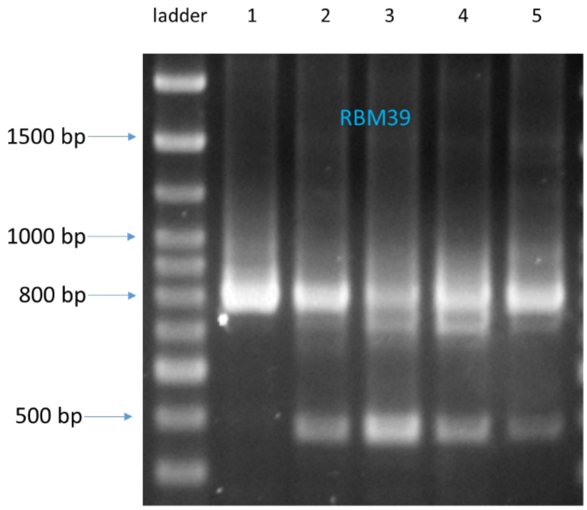


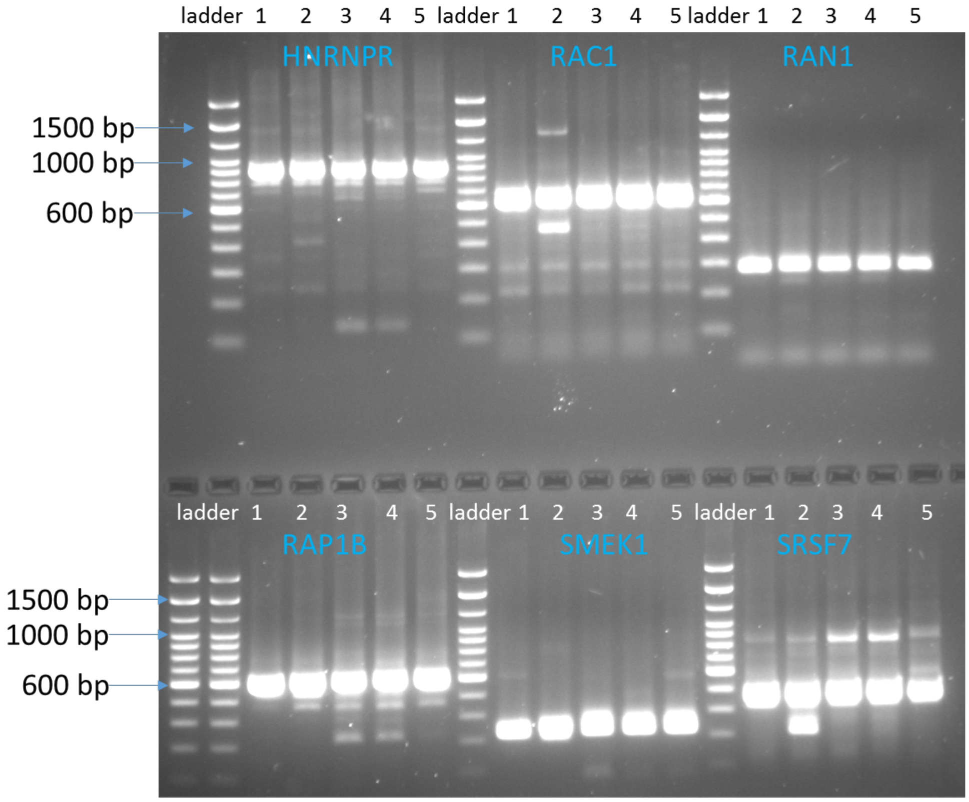

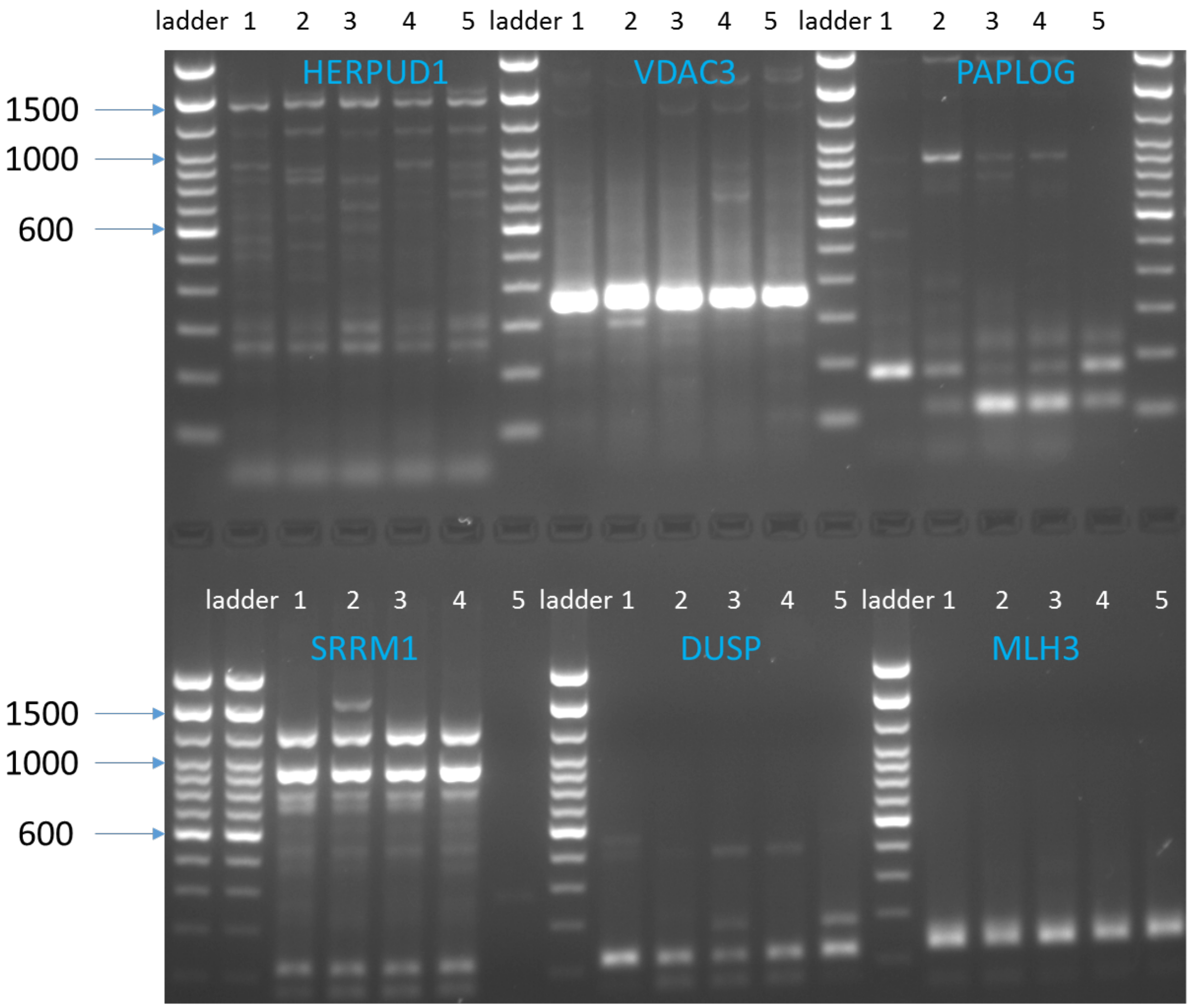


Figure S3. Effects of SD6, **1**, **2**, and **3** on the change in the pre-mRNA splicing for genes using RT-PCR. RT-PCR of 13 genes in RH-18 cells with treatment for 4 h. DNA Ladder; (Lane 1) 0.5% DMSO; (Lane 2) SD6: 10 μM; (Lane 3) **1**: 10 μM; (Lane 4) **2:** 5 μM; (Lane 5) **3**: 10 μM. Clear examples of changes in AS were identified with compound treatments for genes RBM39, HNRNPR, RAP1B, SRSF7, PAPLOG and DUSP.

**E. Tool Compound Dose Response**

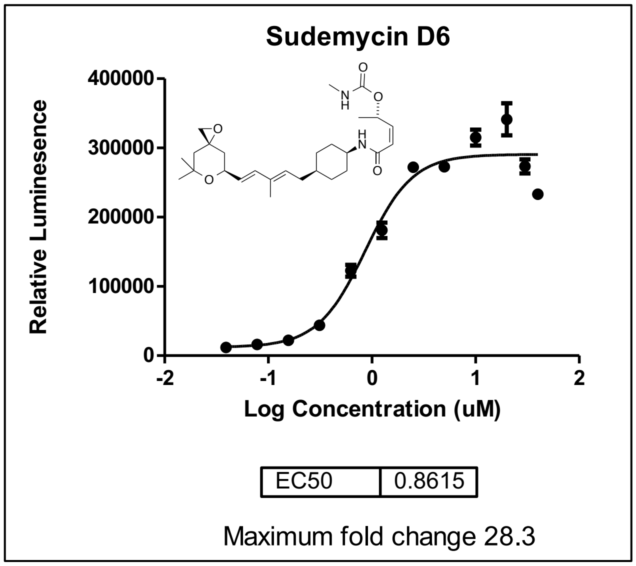


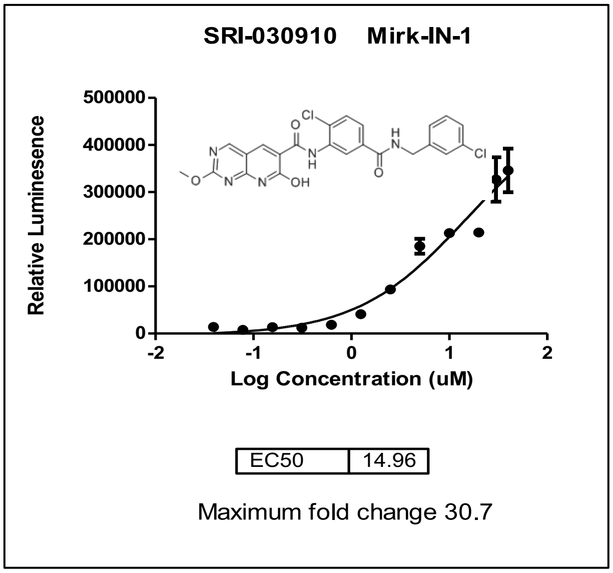

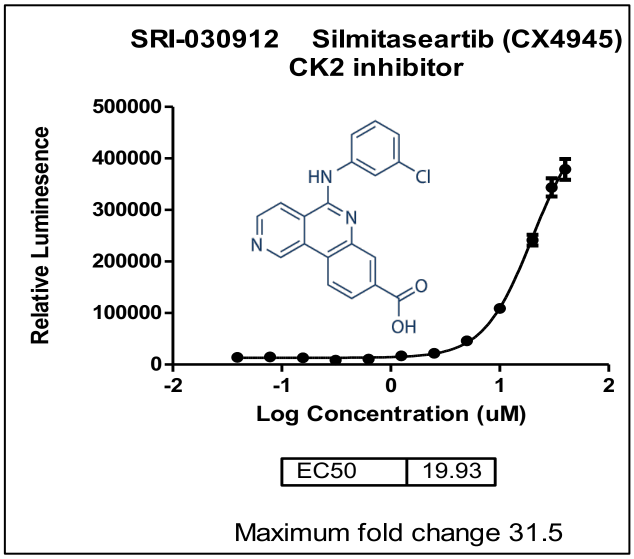


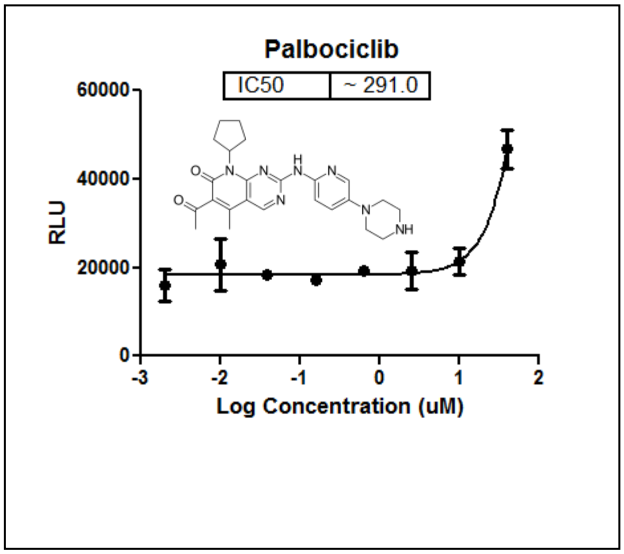


Figure S4. SK-MEL-2/Luc-MDM2 stable cells were plated at a density of 20,000 cells/well in 96-well plates and incubated overnight at 37°C in 5% $CO_2$. The following day, cells were treated with serial dilutions of the indicated compounds for 4 hours and ONE-Glo™ EX reagents (Promega) were added to measure the luciferase activity. Curves not shown for drugs with $IC_{50}$s >> 10 µM.

### F. Antagonist Screen and Assay

#### Antagonist Screen

SK-MEL-2/Luc-MDM2 stable cells were plated at a density of 20,000 cells/well in 96-well plates and incubated overnight at 37°C in 5% $CO_2$.(Shi, Joyner et al. 2015, Shi, Park et al. 2017) The following day, cells ware treated with the 14 analogs (see Table S2 and S3 below) using serial dilution of for one hour. After one-hour pre-treatment, 4 µM of sudemycin D6 were added to the pre-treated cells for another 4 hours. Either 0.5% DMSO or serial dilutions of sudemycin D6 were used to treat cells for 4 hours as controls. At the end of incubation time, ONE-Glo™ EX reagents (Promega) were added to measure the luciferase activity. The graph shows the luc intensity value of DMSO or 4 µM of sudemycin D6 that were the average of the reading of cells treated with DMSO or 4 µM of sudemycin D6 for 4 hours only. The list of compounds screened as potential TESLR antagonists are show in Tables S4 and S5 below. These compounds showed no detectable or weak cytotoxic effects at 10 µM, and so were deemed 'inactive' according to our published standard criteria (72 h cytotox $EC_{50}$ > 1 µM).(Lagisetti, Palacios et al. 2013)

### Antagonist Assay

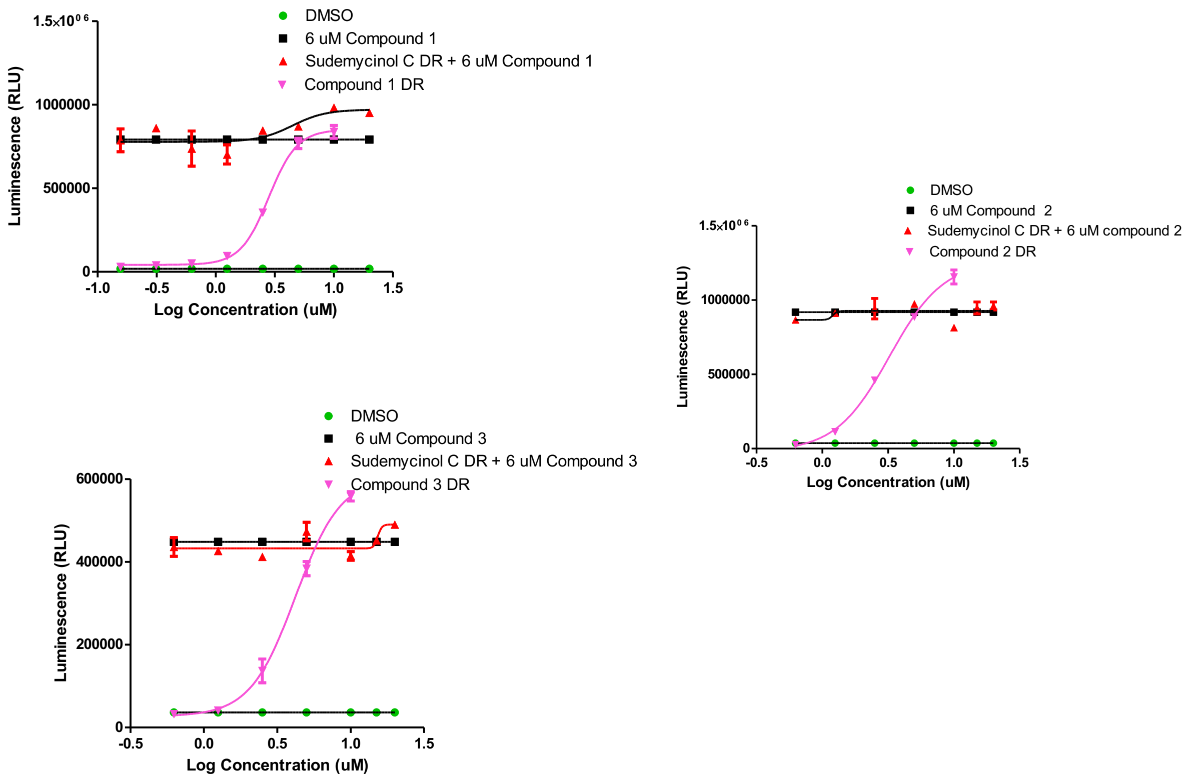


Figure S3. SK-MEL-2/Luc-MDM2 stable cells were plated at a density of 20000/well in 96-well plates and incubated overnight at 37°C in 5% $CO_2$. The following day, some cells were treated with 6 μM of compound **1**, **2** or **3** for one hour. After one-hour pre-treatment, serial dilutions of the sudemycinol C were added to the pre-treated cells for another 4 hours. At the same time, some cells were treated with 0.5% DMSO or serial dilutions of the compound **1**, **2** or **3** only for 4 hours. At the end of incubation time, ONE-Glo™ EX reagents (Promega) were added to measure the luciferase activity. On the graph, value of DMSO and 6 μM of compound **1**, **2** or **3** were the average of the reading.

### G. Immunoblotting Analysis

#### Western Blot Methods

HCT116 cells were seeded at 3 million cells per 100 mm culture plate. The next day cells were treated with 5 µM or 10 µM of each compound for 4h and 8h. For harvesting, cells were washed 2X with ice cold PBS and then resuspended in 400 µl of nuclear isolation lysis buffer (150 mM NaCl, 10 mM Tris-HCl pH 7.4, 1 mM EDTA pH8, 0.5% NP-40, 1X Halt Protease and Phosphatase Inhibitor). Cells were scraped and transferred to a microfuge tube and spun at 2800 rpm for 5 minutes, 4C. The supernatant was removed and the nuclear pellet was resuspended in a nuclear extraction buffer (400 mM NaCl, 20 mM Hepes pH 7.9, 1 mM EDTA pH 8, 1 mM DTT, 1X Halt Protease and Phosphatase Inhibitor). The pellet was snap frozen on dry ice and then thawed and spun at 12K rpm for 5 min, 4C. The supernatant was collected (nuclear extract) and protein concentrations determined using BCA Protein Assay Kit. Samples were run alongside molecular weight markers (Invitrogen SeeBlue™ Plus2 #LC5925) on 4-12% Bis-Tris protein gels (Invitrogen# NP0323BOX) using MOPS SDS running buffer (Invitrogen# NP0001). The proteins were transferred to PVDF membrane using a wet transfer system. The membranes were incubated for 30 minutes at room temperature in Odyssey blocking buffer (Li-Cor 927-40000), then incubated overnight at 4°C with primary antibody in blocking buffer containing 0.2% Tween-20 and 1h at 4°C with secondary antibody in blocking buffer containing 0.2% Tween-20 and 0.02% SDS. After the secondary antibody incubation, membranes were washed 4X with PBS containing 0.1% Tween-20 and 1X with PBS before imaging using the Li-Cor Odyssey imaging system. The primary antibodies used were as follows: SR Monoclonal Antibody (16H3) (Thermo Fisher #16H3E8), Phospho-SF3B1 (Thr313) (D8D8V) (Cell Signaling #25009), Anti-SF3B1 antibody [EPR11987(B)] (Abcam#170854), Anti-phosphoepitope SR protein (clone 1H4) (Millipore# MABE50). The secondary antibodies used are as follows: IRDye® 800CW Goat-anti-Mouse (Li-Cor 925-32210) and IRDye® 680RD Goat-anti-Rabbit (Li-Cor 926-68071).

### H. Natural Product Analog Synthetic Chemistry

Table S4. The list of sudemycin analogs used to screen for TESLR antagonists, the syntheses of these derivatives have been published,(Lagisetti, Pourpak et al. 2008, Lagisetti, Pourpak et al. 2009, Lagisetti, Palacios et al. 2013) except for CLA-G-10. The experimental procedure for this latter compound is described below.

| SRI Number | Compound ID | Structure | MW |
|---|---|---|---|
| SRI030878 | CLA-G-03A | 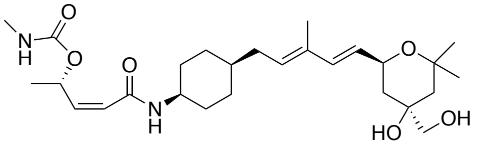 | 492 |
| SRI030879 | CLA-G-03B | 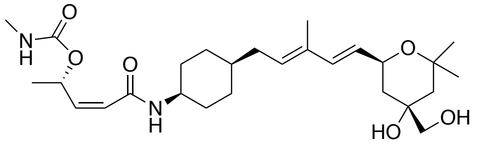 | 492 |
| SRI030880 | CLA-G-10 | 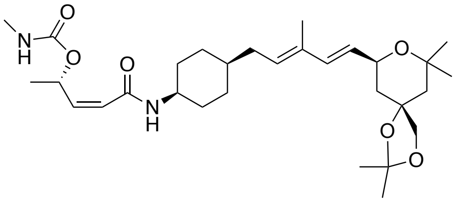 | 532 |
| SRI030881 | CLA-B-69<br>Sudemycinol E | 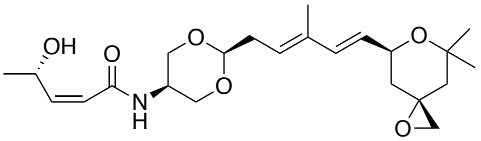 | 421 |
| SRI030882 | 16618-CL-43<br>Sudemycinol C | 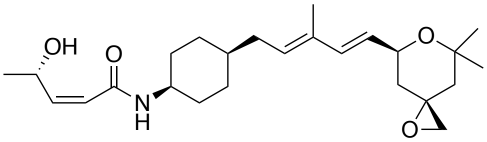 | 417 |
| SRI030883 | CLA-D-42 | 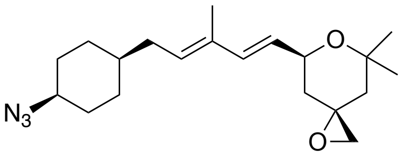 | 345 |
| SRI030884 | CLA-B-96 | 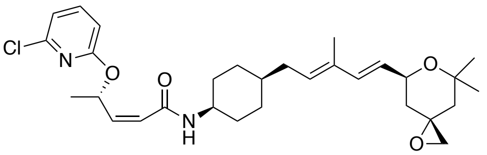 | 529 |

| | | | |
|---|---|---|---|
| SRI030885 | CLA-C-03 | 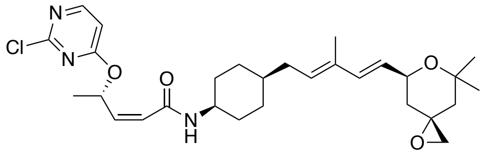 | 530 |
| SRI-030886 | CLA-C-06 | 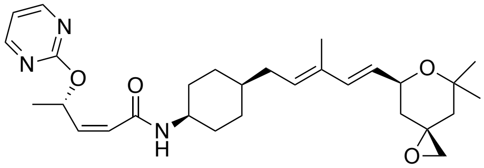 | 495 |

**Table S5**. The list of herboxidiene analogs used to screen for TESLR antagonists, the synthesis of these derivatives has been published,(Lagisetti, Yermolina et al. 2014) except for CLA-E-39 and CLA-E-40. The experimental procedure for the new compounds is described below. Abbreviations: DEIPS= diethylisopropylsilyl, Me= methyl.

| **SRI Number** | **Compound ID** | **Structure** | **MW** |
|---|---|---|---|
| SRI030887 | CLA-E-39 | 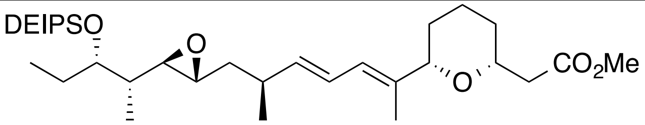 | 494 |
| SRI030888 | CLA-E-40 | 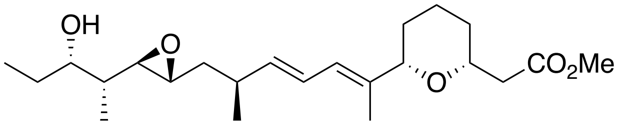 | 394 |
| SRI030889 | CLA-E-72 | 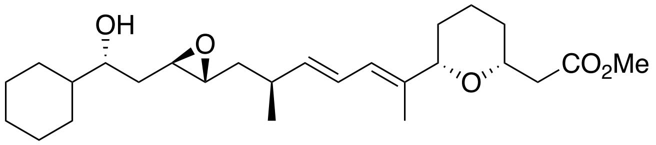 | 448 |
| SRI030890 | CLA-E-59 | 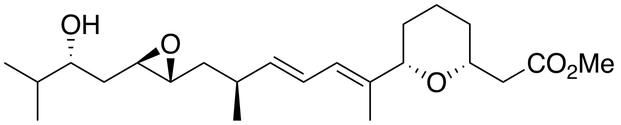 | 394 |
| SRI030891 | CLA-E-91 | 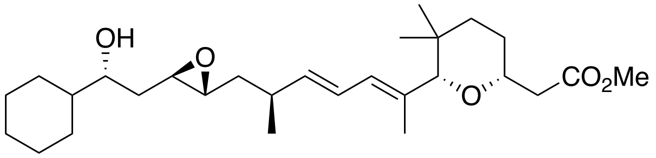 | 476 |

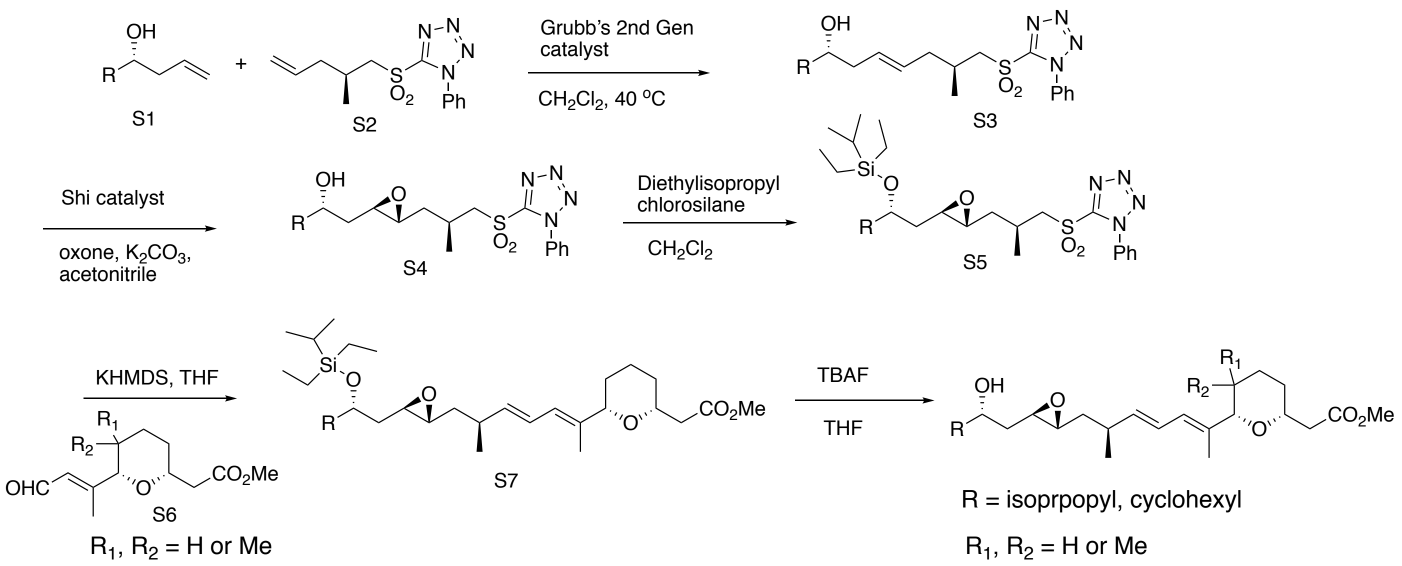

Scheme S1. General scheme for the synthesis of herboxidiene analogs.

**Synthetic Procedures for Natural Product Analogs**

The following compounds have not been previously reported but were prepared based on our published work.(Lagisetti, Pourpak et al. 2008, Lagisetti, Pourpak et al. 2009, Lagisetti, Palacios et al. 2013, Lagisetti, Yermolina et al. 2014) All of the other compounds have been published as reported above.

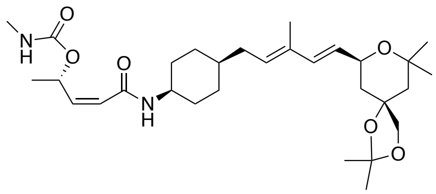

**CLA-G-10:** CLA-G-10 is prepared from diol CLA-G-03B by treatment with 2,2-dimethoxy propane and catalytic toluene-4-sulfonic acid. $^{1}$H NMR (400 MHz, Chloroform-*d*) δ 6.17 (d, *J* = 15.7 Hz, 1H), 5.75 (d, *J* = 11.4 Hz, 1H), 5.59 - 5.33 (m, 3H), 4.31 (dd, *J* = 11.3, 6.7 Hz, 1H), 3.67 - 3.52 (m, 2H), 2.67 (d, *J* = 4.8 Hz, 3H), 1.99 (t, *J* = 7.1 Hz, 2H), 1.72 - 1.41 (m, 17H), 1.29 (d, *J* = 9.0 Hz, 12H), 1.22 - 1.18 (m, 3H), 1.15 - 1.11 (m, 3H); $^{13}$C NMR (101 MHz, $CDCl_3$) δ 165.15, 157.04, 135.85, 133.84, 131.77, 127.33, 126.14, 109.93, 78.69, 75.51, 72.61, 69.18, 68.43, 45.33, 44.88, 41.51, 36.69, 32.57, 30.97, 29.59, 29.46, 27.66, 27.44, 27.40, 27.31, 27.21, 25.83, 24.38, 20.62, 12.46; MS (ESI) *m/z* 533 $(M+1)^+$

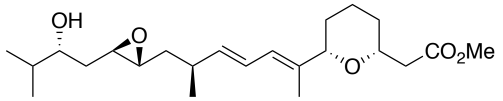

**CLA_E_59:** $^{1}$H NMR (400 MHz, Chloroform-*d*) δ 5.99 (dt, *J* = 15.7, 0.9 Hz, 1H), 5.48 (dd, *J* = 15.7, 8.1 Hz, 1H), 5.30 (d, *J* = 7.5 Hz, 1H), 4.17 - 4.02 (m, 1H), 3.86 - 3.70 (m, 1H), 3.60 (s, 3H), 3.50 - 3.40 (m, 1H), 2.84 - 2.70 (m, 2H), 2.58 - 2.45 (m, 1H), 2.40 - 2.27 (m, 2H), 1.88 (d, *J* = 4.5 Hz, 1H), 1.84 - 1.77 (m, 1H), 1.68 – 1.61 (m, 1H), 1.74 - 1.68 (m, 4H), 1.62 - 1.51 (m, 4H), 1.38 - 1.15 (m, 3H), 1.02 – 0.96 (m, 3H), 0.87 – 0.81 (m, 6H); $^{13}$C NMR (101 MHz, $CDCl_3$) δ 171.83, 135.92, 134.30, 133.36, 130.96, 74.95, 74.20, 73.92, 57.41, 57.33, 51.58, 41.48, 39.71, 35.42, 35.10, 33.77, 31.31, 30.80, 23.38, 21.21, 18.56, 17.40, 13.13; MS (ESI) *m/z* 395 $(M+1)^+$

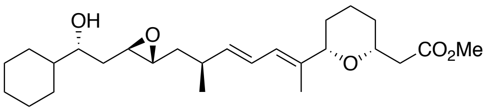


**CLA_E_72**: $^1$H NMR (400 MHz, Chloroform-*d*) δ 6.27 (ddt, *J* = 14.2, 10.9, 1.6 Hz, 1H), 6.01 (d, 14.2 Hz, 1H), 5.60 - 5.51 (m, 1H), 3.83 (ddd, *J* = 11.4, 7.6, 5.7 Hz, 1H), 3.73 (d, *J* = 11.0 Hz, 1H), 3.68 (s, 1H), 3.50 (d, *J* = 10.0 Hz, 1H), 2.93 - 2.76 (m, 2H), 2.64 - 2.56 (m, 1H), 2.50 - 2.39 (m, 2H), 1.95 (s, 1H), 1.89 (dt, *J* = 12.9, 2.9 Hz, 1H), 1.81 - 1.71 (m, 7H), 1.70 - 1.60 (m, 6H), 1.59 - 1.50 (m, 4H), 1.41 - 1.14 (m, 7H), 1.10 - 1.05 (m, 3H), 1.04 - 0.95 (m, 2H); $^{13}$C NMR (101 MHz, $CDCl_3$) δ 171.85, 138.97, 136.92, 125.27, 124.13, 82.09, 74.44, 73.38, 57.37, 57.35, 51.56, 43.76, 41.51, 39.70, 35.49, 35.30, 31.06, 29.61, 29.01, 27.93, 26.49, 26.23, 26.10, 23.55, 21.12, 13.60; MS (ESI) *m/z* 449 $(M+1)^+$

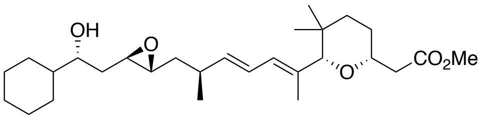


**CLA_E_91:** $^1$H NMR (400 MHz, Chloroform-*d*) δ 6.22 (dd, *J* = 15.3.0, 10.9 Hz, 1H), 5.85 (dd, *J* = 11.0, 1.6 Hz, 1H), 5.51 (ddd, *J* = 15.3, 8.4, 2.8 Hz, 1H), 3.81 - 3.71 (m, 1H), 3.67 (s, 3H), 3.52 (d, *J* = 10.7 Hz, 2H), 2.92 - 2.78 (m, 2H), 2.68 - 2.58 (m, 1H), 2.46 (ddd, *J* = 14.9, 6.8, 4.8 Hz, 2H), 1.98 (t, *J* = 4.8 Hz, 1H), 1.85 – 1.71 (m, 8H), 1.70 - 1.59 (m, 3H), 1.56 - 1.45 (m, 6H), 1.36 -1.29 (m, 1H), 1.25 - 1.12 (m, 3H), 1.07 (d, *J* = 6.8 Hz, 3H), 1.04 - 0.92 (m, 2H), 0.88 (s, 3H), 0.82 (s, 3H); $^{13}$C NMR (101 MHz, $CDCl_3$) δ 171.84, 138.52, 134.73, 127.34, 125.13, 89.99, 74.62, 73.36, 57.47, 57.43, 51.54, 43.78, 41.35, 39.71, 39.60, 35.44, 35.30, 33.84, 29.00, 28.16, 28.01, 27.96, 26.48, 26.24, 26.10, 21.16, 20.42, 15.01; MS (ESI) *m/z* 477 $(M+1)^+$
